# Localizing genomic regions contributing to the extremes of externalizing behavior: ADHD, aggressive and antisocial behaviors

**DOI:** 10.1101/750091

**Authors:** Mariana L. Rodríguez-López, Hilleke Hulshoff Pol, Barbara Franke, Marieke Klein

**Author notes:** Corresponding author: Marieke Klein | | UC San Diego, Department of Psychiatry | 9500 Gilman Dr, La Jolla, CA 92093-0667, US.

## Abstract

Attention-Deficit/Hyperactivity Disorder (ADHD) is a neurodevelopmental disorder, which in some cases occurs comorbid with aggressive and antisocial behavior (AGG; ASB). The three externalizing behaviors are moderately to highly heritable and are genetically correlated. However, the genomic regions underlying this correlation are unknown. In this study, we aimed to localize genetic loci shared between ADHD, AGG, and ASB, using two complementary approaches.

GWAS summary statistics for ADHD, AGG, and ASB were used for (1) cross-trait gene-based meta-analysis association analyses and (2) local genetic correlation analyses to identify shared genetic loci. Results of both complementary methods were combined to retrieve overlapping genes. Biological functionality of prioritized genes was assessed by exploring gene expression patterns in brain tissues and testing for gene-based association with (subcortical) brain regions.

We confirmed previous findings that ADHD, AGG, and ASB were positively genetically correlated at a global level. We identified eleven significant genes in cross-trait gene-based meta-analyses, 31 loci shared between traits; 34 genes were identified when both approaches were combined.

This study emphasizes the complex genetic architecture underlying global genetic correlations at the locus level. Converging evidence from these cross-trait analyses highlights novel candidate genes underlying biological mechanisms shared by ADHD, AGG, and ASB.

## Introduction

Attention-deficit/hyperactivity disorder (ADHD) is a common neurodevelopmental disorder with a world-wide prevalence of 5% in children and 2.5% in adults (Polanczyk, de Lima, Horta, Biederman, & Rohde, 2007; Simon, Czobor, Bálint, Mészáros, & Bitter, 2009). It is characterized by developmentally inappropriate and impairing levels of inattention and/or hyperactivity and impulsivity. Twin and family studies revealed that ADHD is highly heritable, with heritability estimates around 74% (Faraone & Larsson, 2019). ADHD’s genetic architecture is characterized by a multifactorial pattern of inheritance, meaning that multiple genetic and environmental factors, most of small effect, and their interplay can contribute to its pathophysiology. A recent genome-wide association study (GWAS) meta-analysis identified the first genome-wide significant associations for ADHD, and the heritability estimated from single nucleotide polymorphisms (SNPs, SNP-based heritability) was 22% (Demontis et al., 2019). Comorbidity is the rule rather than the exception in ADHD, and individuals with ADHD often also show co-occurring psychiatric and behavioral disorders and traits, such as autism spectrum disorder, schizophrenia, and major depressive disorder. Aggressive behavior (AGG) and disruptive behavioral disorders as well as antisocial behavior (ASB) and antisocial personality disorder co-occur frequently with ADHD, as different studies showed high prevalence of comorbid ADHD with Oppositional Defiant Disorder (ODD)/Conduct Disorder (CD), ranging from 25% up to 80% (reviewed in (Franke et al., 2018)). Importantly, the phenotypic and genetic overlap between externalizing behaviors and related disorders is not expected to be complete and these phenotypes also exist while not overlapping phenotypically.

AGG is defined as the intended act (physical, verbal, or psychological) of causing harm to others or to oneself (Liu, 2004). ASB refers to actions and attitudes that affect others and violate societal norms (Burt & Neiderhiser, 2009; Lahey et al., 2008). ASB includes behaviors with an aggressive connotation, like bullying or mugging, but also includes non-physically aggressive behaviors like stealing, truancy, and lying (Lahey et al., 2008). Family and twin studies showed that both AGG and ASB are moderately to highly heritable (~50-70%) (Ferguson, 2010; Hudziak et al., 2003; Miles & Carey, 1997; Rhee & Waldman, 2002). So far, one GWAS meta-analysis on childhood AGG and one for ASB has been published, both including individuals from the general population (Pappa et al., 2016; Tielbeek et al., 2017). For both studies, the obtained SNP-based heritability estimates were low (h2~5%).

The genetic mechanisms underlying the co-occurrence of ADHD and AGG and ASB have been investigated through family and twin studies. Several studies suggested that ADHD with AGG or ASB could be a separate subtype of ADHD, based on the different development, persistence, and risk-seeking seen in those individuals showing the combined phenotypes (Christiansen et al., 2008; Doyle & Faraone, 2002; Retz & Rösler, 2009). Results of molecular genetic studies of common variants showed positive and significant genetic correlations of ADHD with AGG and ASB (ADHD-AGG rg=~0.7 and ADHD-ASB rg=~0.55) (Zhang-James et al., 2019). Individuals with co-occurring symptoms were found to have a higher ADHD persistence rate and overall higher clinical severity, pointing out that the relationship between the disorder and traits is related to a greater load of shared genetic factors (Hamshere et al., 2013; Zhang-James et al., 2019).

Global approaches assessing the genetic correlations between two traits/disorders may over-generalize the relationship between them and may lack specificity in terms of the actual causal genes involved in shared genetic etiology. In this study, using GWAS summary statistics, we aimed to localize and define the contribution of shared genetic loci between ADHD, AGG, and ASB by combining two complementary approaches (study flow schematically shown in **Figure 1**). Identified loci can give us insights into the genetic complexity underlying the extremes of externalizing behavior. Finally, we localized genetic effects in the brain and explored gene expression patterns in relevant brain tissues and prioritized candidate genes for future, in depth studies.

**Figure 1.**
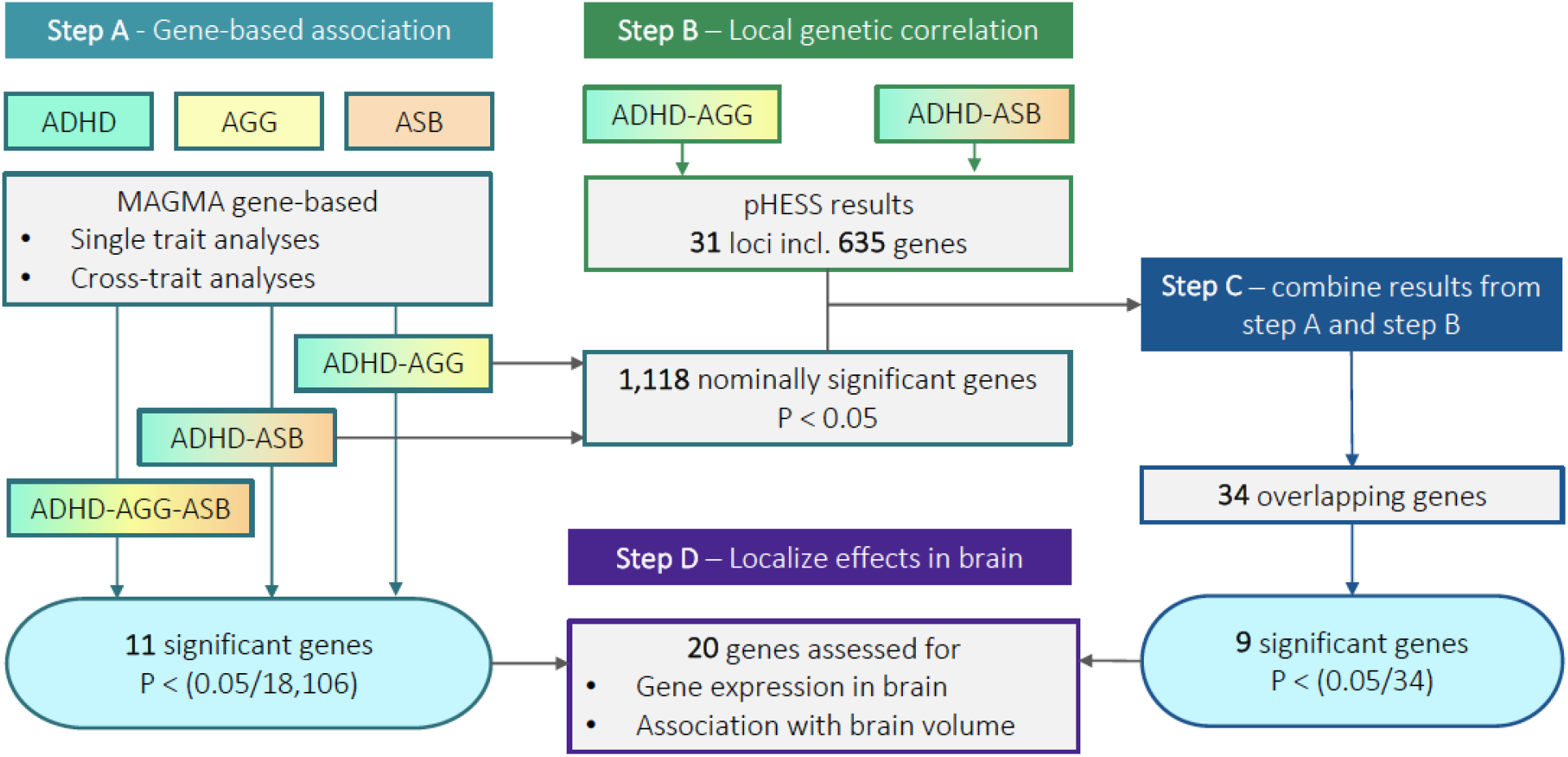
Schematic overview of study flow to identify overlapping genes between ADHD, AGG, and ASB. **Step A)** We performed single trait gene-based association analyses by using MAGMA (https://ctg.cncr.nl/software/magma) for each trait separately, followed by three cross-trait gene-based meta-analyses. Eleven genes showed significant gene-based association. All genes with a gene-based p-value < 0.05 were selected for Step C). **Step B)** Local genetic correlation analyses were performed for ADHD+AGG and ADHD+ASB. Genes were mapped to 31 loci by uploading loci positions to the UCSC Genome Browser (https://genome.ucsc.edu). **Step C)** We assessed the overlapping genes between the two methods. Out of those 34 genes, we selected nine genes that showed nominal gene-based association with brain volumes using ENIGMA data (p< 0.05/34). **Step D)** Localizing the effects in the brain was completed for 20 genes, which were analyzed for expression in brain tissue and association with (subcortical) brain volume.

## Methods

### Samples

In this study, we used GWAS summary statistics (GWAS-ss) data obtained from publicly available independent meta-analysis studies of ADHD, AGG, and ASB. These studies had been approved by local ethics committees and had obtained the required informed consents (as described in earlier publications (Demontis et al., 2019; Pappa et al., 2016; Tielbeek et al., 2017)). All data had to meet the following criteria: 1) participants must be of European ancestry and 2) variants had to be annotated in human genome assembly hg19. Moreover, only data of autosomal chromosomes were considered. Because of its unusual LD structure, we removed the Major Histocompatibility Complex (MHC) region from the analyses (chr6:28477797-33448354). Further information on the primary GWAS meta-analyses is provided in <u>supplementary material</u> or in their original (Demontis et al., 2019; Pappa et al., 2016; Tielbeek et al., 2017). A short summary with the main characteristics of each of these studies can be found below.

The ADHD data was derived from 19,099 cases and 34,194 controls including 12 cohorts of the Psychiatric Genomics Consortium (PGC) and the Lundbeck Foundation Initiative for Integrative Psychiatric Research (iPSYCH) ((Demontis et al., 2019), https://www.med.unc.edu/pgc/download-results/).

The GWAS meta-analysis data for aggressive behavior (AGG) was provided by the study by Pappa and colleagues (Pappa et al., 2016), performed in the framework of the Early Genetics and Lifecourse Epidemiology (EAGLE) consortium (http://www.tweelingenregister.org/EAGLE/). The authors had meta-analyzed data from nine cohorts including children between the age of 3 to 15 years, with a total of 18,988 participants.

For antisocial behavior (ASB), we used the summary statistics from a study of the Broad Antisocial Behavior Consortium (BroadABC; http://broadabc.ctglab.nl/summary_statistics) (Tielbeek et al., 2017)). The meta-analysis consisted of five discovery cohorts of European ancestry, including 16,400 individuals in total (mean age 6.7 – 43.8 years).

GWAS meta-analysis summary statistics for intracranial volume (ICV), and seven subcortical brain volumes (nucleus accumbens, amygdala, caudate nucleus, hippocampus, pallidum, putamen, and thalamus) were obtained from the combined datasets of the Enhancing Neuro Imaging Genetics through Meta-Analysis (ENIGMA) consortium and the Cohorts for Heart and Aging Research in Genomic Epidemiology (CHARGE) consortium including almost 40,000 European ancestry participants, (Adams et al., 2016; Hibar et al., 2017; Satizabal et al., 2019); unrestricted data, not including the replication and generalization samples, was requested and downloaded from http://enigma.ini.usc.edu/research/download-enigma-gwas-results/.

### Genome-wide genetic correlation analyses

We conducted linkage disequilibrium score regression analysis (LDSC) (Bulik-Sullivan et al., 2015) using the latest ADHD GWAS to (re-)estimate the genetic correlation between ADHD and AGG, and between ADHD and ASB. Data processing in LDSC included an automatic QC step and the use of pre-computed LD score estimates based on European samples from the 1000 Genomes (Abecasis et al., 2012), both indicated in https://github.com/bulik/ldsc/wiki/Heritability-and-Genetic-Correlation.

### Gene-based analyses

Gene-based association analyses were performed using the MAGMA software (version 1.05; (de Leeuw, Mooij, Heskes, & Posthuma, 2015)). SNPs were mapped onto genes and then the association (computing gene-based p-values) of each gene with a phenotype of interest (ADHD, AGG, and ASB and the eight brain volumes [nucleus accumbens, amygdala, caudate nucleus, hippocampus, ICV, pallidum, putamen, thalamus]) was tested using the summary statistics from the SNP-based GWAS meta-analyses **(Figure 1, step A,D)**. By using the weighted Stouffer’s Z method as implemented in MAGMA, cross-trait gene-based meta-analyses were performed, resulting in gene-based association p-values for ADHD-AGG, ADHD-ASB, and ADHD-AGG-ASB (**Figure 1, step A**). For subsequent analyses, we prioritized genes that (1) had a statistically significant p-value in the cross-trait meta-analyses results, (2) had a smaller p-value in the cross-trait meta-analyses results compared to the gene-based p-values of the individual traits, and (3) had at least a nominally significant gene-based p-value in the individual trait results. For the gene-based meta-analyses including ADHD, AGG, and ASB data, the Bonferroni-corrected significance threshold was set to p < 2.8e-06 (0.05/18,106 genes).

### Local genetic correlation analyses

To assess which genomic regions were contributing most to the genome-wide SNP-based heritability estimates of the traits of interest, we used the software package HESS version 0.5.4-beta (Shi, Kichaev, & Pasaniuc, 2016). Working with GWAS-ss, HESS estimates for each one of the loci defined in a genome partition file the variance explained by all the typed SNPs at a single locus while taking into account the LD pattern between the SNPs. While traditional models assessing the global SNP-based heritability assume that all SNPs contribute to the trait (and this model is invalid at most risk loci), HESS uses a more robust approach as it does not assume any distribution for the effect sizes at causal variants. We used a genome partition file from a previous study (Berisa & Pickrell, 2016) and a SNP reference file from 1000 Genomes Project for the European population panel (Abecasis et al., 2012) to define a total of 1,694 LD-independent loci. Both files can be downloaded directly from https://huwenboshi.github.io/hess/input_format/. Its extension, ρ-HESS (Shi, Mancuso, Spendlove, & Pasaniuc, 2017), was used to obtain cross-trait genetic covariance estimates on genomic regions of the traits of interest. We calculated the local genetic correlation, the standardized version of the local genetic covariance, for each of the 1,694 loci for both cross-trait analyses across the two cross-trait analyses (ADHD-AGG and ADHD-ASB). The calculation looked similar to the one described by Bulik-Sullivan and colleagues for genome-wide genetic correlation (Bulik-Sullivan et al., 2015), but was modified by Shi and colleagues to obtain the genetic correlation for the *i*th region (rg,local,*i*) by using both local heritability estimate of trait one (SNPh1^2^local,*i*) and trait two (SNPh2^2^local,*i*) and local genetic covariance (ρg,local,*i*) as shown in **Equation 1** (Shi et al., 2017).

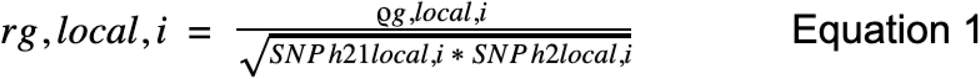

SNP-based heritability of regions was considered to be significant at a Bonferroni-corrected p-value threshold of p < 2.95e-05 (0.05/1,694). We only considered loci with a local genetic correlation score between minus one and one [-1,1], which are the expected values for a correlation analysis. Next, we selected the loci that were overlapping between the ADHD-AGG and ADHD-ASB cross-trait analyses. Finally, by using the UCSC Genome Browser, we identified the genes located in the selected loci for the two cross-trait pairs (**Figure 1, step B**).

### Combining results of MAGMA and ρ-HESS analyses

We combined results of MAGMA and ρ-HESS analyses and selected all overlapping genes with a nominally significant gene-based association p-value for each cross-trait gene-based meta-analysis (**Figure 1, step C**). Next, we selected overlapping genes which had a significant p-value (Bonferroni corrected threshold [p < 0.05/number of genes tested], **step C**), for subsequent biological annotations (**step D**) in at least one of the cross-trait analyses.

### Biological annotation of candidate genes

To retrieve more information about relevant biological mechanisms and to prioritize the selected set of genes, we used gene expression analysis and tested for association with brain volumes.

First, to assess mRNA expression profiles of our candidate genes, we used RNA-sequencing data from 54 tissues, provided by Genotype-Tissue Expression (GTEx) Project V8 ((Battle, Brown, Engelhardt, & Montgomery, 2017), https://gtexportal.org/home/). For each gene, we obtained the median Transcripts Per Million (TPM) in each tissue. We prioritized those brain tissues that belong to the top 5% (3 out of all 54 tissues) that most strongly express the gene. Among all 54 tissues, we present results on the 12 available brain tissues: amygdala, anterior cingulate cortex, caudate nucleus, cerebellar hemisphere, cerebellum, cortex, frontal cortex, hippocampus, hypothalamus, nucleus accumbens, putamen, and substantia nigra (Battle et al., 2017). Second, we performed gene-based association analyses (as described in the “gene-based analysis” section) using data for the volumes of seven brain regions (nucleus accumbens, amygdala, caudate nucleus, hippocampus, pallidum, putamen, and thalamus) and for intracranial volume (ICV) (Adams et al., 2016; Hibar et al., 2017; Satizabal et al., 2019). Genes were considered significant with a Bonferroni-corrected p-value threshold of p < 3.13e-04 (0.05/160 [number of genes tested x number of tissues]).

## Results

### Genome-wide genetic correlation analyses

We assessed and confirmed global genetic correlations between ADHD, AGG, and ASB. The genetic correlation between ADHD and AGG was rg = 0.7 (SE = 0.18, p = 0.0001), the one for ADHD and ASB was rg = 0.51 (SE = 0.19, p = 0.006), as published previously (Zhang-James et al., 2019). The genetic correlation between AGG and ASB was lower and non-significant (rg = 0.24, SE = 0.36, p = 0.503).

### Localizing shared genetic loci

First, we performed three cross-trait gene-based meta-analyses and selected all genes with a significant gene-based p-value, which showed stronger association in cross-trait meta-analyses compared with the individual trait analyses (**Figure 1, step A**). In the ADHD+AGG analysis, we identified two significant genes *(ZNF521* and *MECOM,* **Table 1**), for ADHD+ASB, we found eight genes to be significant *(SZT2, HYI, DDN, FAM186B, DUSP6, GPM6A, COL19A1,* and *LY96,* **Table 1**), and for the analysis of ADHD+AGG+ASB, four genes were significant (*HYI*, *GALNT3, MECOM,* and *LY96,* **Table 1**). *GALNT3* was the only candidate that specifically emerged in the three-trait gene-based meta-analysis. In our second approach, we estimated which regions contributed most to the phenotypic variance of a trait explained by the SNPs in 1,694 LD-independent loci. We used ρ-HESS to analyze the ADHD+AGG and ADHD+ASB trait pairs and estimated the local genetic covariance for these loci. We observed varying local genetic correlation estimates throughout the genome, including both positively and negatively correlated loci with some having higher genetic correlation scores than others, contrary to the usually expected equal contribution from all loci (**Figure 2**). Among the regions were those for which both associations, ADHD+AGG and ADHD+ASB, appeared to have a positive local genetic correlation (e.g. chr4:6,773,043-7,539,692; ADHD-AGG rg = 0.71, ADHD-ASB rg = 0.6), while other loci seemed to show a positive correlation in one and a negative correlation in the other analysis (e.g. chr1:196,176,201-197,311,514; ADHD-AGG rg = 0.99, ADHD-ASB rg = −0.96), and for others negative genetic correlations for both associations were observed (e.g. chr12:124,977,980-126,445,505; ADHD-AGG rg = −0.13, ADHD-ASB rg = −0.53, **Supplementary Table 1**). As we did not identify significant loci after correction for multiple testing for either of the ρ-HESS-based approaches, we focused all subsequent analyses on the 31 loci (including 635 genes) which overlapped between both ρ-HESS-based cross-trait analyses.

**Figure 2.**
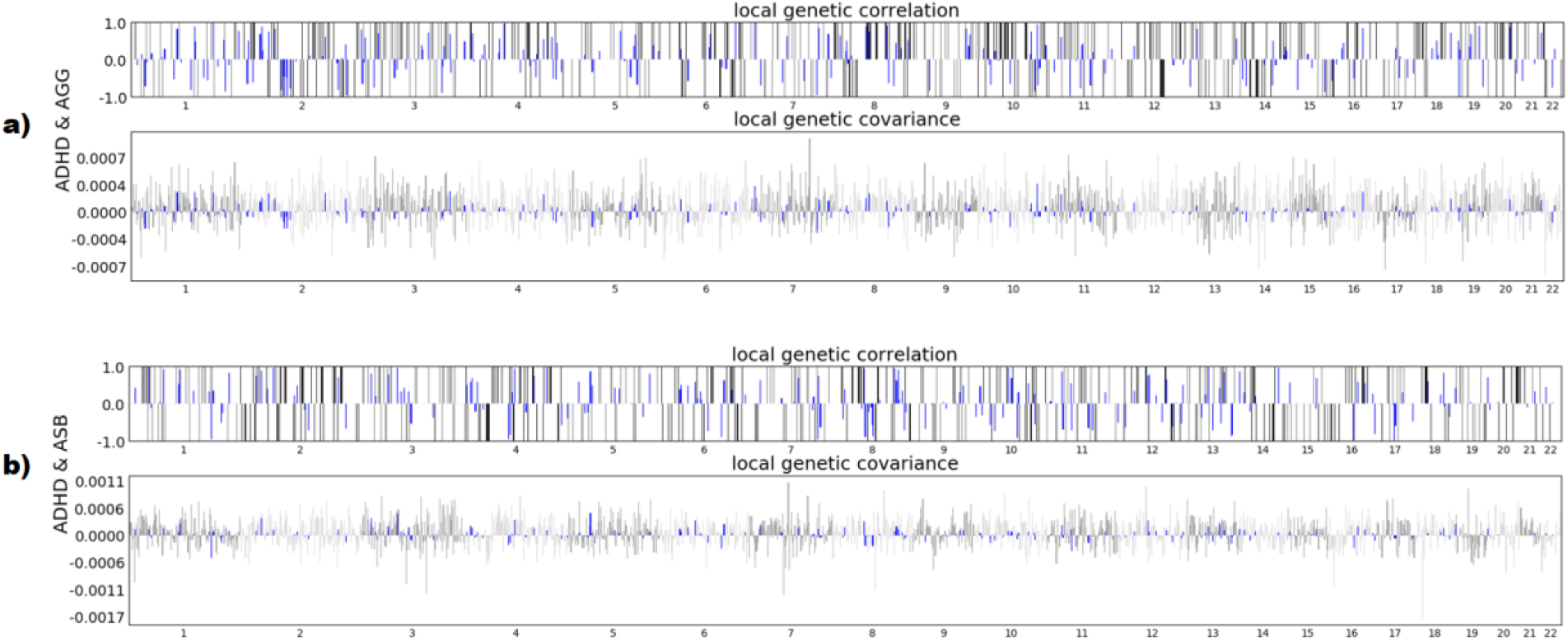
Bar plots of local genetic covariance and correlation analysis using ρHESS for ADHD+AGG and ADHD+ASB cross-trait analyses. a) Local genetic covariances plotted for ADHD+AGG cross-trait analysis. b) Local genetic covariances plotted for ADHD+ASB cross-trait analysis. For both a) and b) the x-axis depicts the loci’s genomic position. The upper bar plot shows the local genetic correlation estimates and the lower bar plots shows the local genetic covariance estimates. Blue bars represent the loci that have expected correlation values inside the −1 to 1 range and which were used for subsequent analyses. Different grey tones distinguish between even and odd chromosome numbers.

**Table 1.** Results of single- and cross-trait gene-based meta-analysis association analyses using MAGMA.

| **A. ADHD and AGG** |  |  |  |
| --- | --- | --- | --- |
| **Gene** | **ADHD gene-based P** | **AGG gene-based P** | **ADHD+AGG MA gene-based P** |
| ***ZNF521*** | 0.000337 | 0.000454 | 2.14e-06 |
| ***MECOM*** | 0.000265 | 2.63e-07 | 1.96e-08 |
| **B. ADHD and ASB** |  |  |  |
| **Gene** | **ADHD gene-based P** | **ASB gene-based P** | **ADHD+ASB MA gene-based P** |
| ***SZT2*** | 1.73e-08 | 0.02544 | 3.77e-09 |
| ***HYI*** | 5.34e-07 | 0.00126 | 5.06e-09 |
| ***DDN*** | 6.65e-05 | 0.00358 | 1.73e-06 |
| ***FAM186B*** | 6.94e-06 | 0.05793 | 2.46e-06 |
| ***DUSP6*** | 5.45e-09 | 0.02422 | 1.23e-09 |
| ***GPM6A*** | 6.24e-05 | 0.00056 | 4.17e-07 |
| ***COL19A1*** | 1.17e-05 | 0.01945 | 1.28e-06 |
| ***LY96*** | 0.00010492 | 0.00010 | 2.46e-07 |
| **C. ADHD, AGG and ASB cross-trait gene-based meta-analysis** |  |  |  |
| **Gene** | **ADHD gene-based P** | **ASB gene-based P** | **ADHD+ASB MA gene-based P** |
| ***HYI*** | 5.34e-07 | 0.50543 | 0.00126 |
| ***GALNT3*** | 0.001399 | 0.011891 | 0.0013519 |
| ***MECOM*** | 0.000265 | 2.63e-07 | 0.37857 |
| ***LY96*** | 0.000105 | 0.16356 | 0.000104 |

### Combining results of MAGMA and ρ-HESS analyses

In combining the most significant results from our two complementary analysis approaches, we derived 34 genes from 31 loci of interest (**Supplementary Table 2**). For this, we selected from the overlapping ρ-HESS-derived loci the annotated genes, in total 635 genes. Among those, we identified genes that also showed (at least) nominally significant association in the gene-based cross-trait analyses (N genes = 1,118; ADHD+AGG or ADHD+ASB, schematically shown in **Figure 1, step C**). Out of the 34 overlapping genes, nine genes were significantly associated with ADHD, AGG and/or ASB in the gene-based analyses (**Figure 1, step C; Table 2**). In total, 11 genes were prioritized from the gene-based association analyses (step A, **Table 1**) and nine genes from combining the MAGMA and ρ-HESS results (step C, **Table 2**).

**Table 2.** Results of single- and cross-trait gene-based meta-analysis association analyses for the prioritized genes by combining ρHESS and MAGMA analyses.

| Gene | ADHD+AGG MA P | ADHD+ASB MA P | ADHD P | AGG P | ASB P |
| --- | --- | --- | --- | --- | --- |
| *PYGO2* | 0,0006* | 0,0004* | 0,0006* | NA | 0,126 |
| *ZBTB41* | 0,0074 | 0,0010* | 0,037 | 0,037 | 0,0007* |
| *ATRNL1* | 0,00002* | 0,0002* | 0,0001* | 0,030 | 0,214 |
| *TBC1D14* | 0,0023 | 0,0003* | 0,0053 | 0,108 | 0,0077 |
| *C5orf51* | 0,0004* | 0,022 | 0,0055 | 0,0087 | 0,675 |
| *FBXO4* | 0,0003* | 0,0097 | 0,0014* | 0,048 | 0,720 |
| *CCDC152* | 0,0014* | 0,0034 | 0,0098 | 0,026 | 0,085 |
| *SEPP1* | 0,0009* | 0,012 | 0,014 | 0,0069 | 0,237 |
| *NKAIN2* | 0,012 | 0,00147* | 0,015 | 0,227 | 0,013 |

### Localize the effects of candidate genes in the brain

Next, we examined the biological characteristics of the 20 identified genes (11 genes identified in **step A** and 9 in **step C**) to localize their effects in the brain in order to assess which brain regions are most frequently implicated in externalizing behaviors. For this, we retrieved information from two different sources for each gene: (1) we assessed whether genes are expressed in relevant brain regions using the GTEx database, and (2) we tested if genes showed significant gene-based association with relevant brain measures from region of interest analyses of subcortical brain volumes by the ENIGMA consortium. Except for *HYI,* all genes were found to be expressed in all 12 brain regions (**Supplementary Table 3**). The cerebellum hemisphere, cerebellum, and frontal cortex (Brodmann Area 9, BA9) showed the highest expression of multiple prioritized genes (**Supplementary Table 3**). Using the ENIGMA data, none of the 20 genes was significantly associated with subcortical brain volume. Twelve genes showed nominally significant associations, and amygdala and putamen were the structures most frequently involved in those associations (**Supplementary Table 4**).

Integrating the findings across the two sources of data, out of the 20 candidate genes, four genes *(ZNF521, DDN, GPM6A,* and *ZBTB41)* are both, highly expressed in brain tissue and showing nominal gene-based association with subcortical brain volumes (**Supplementary Tables 3 & 4**). We also prioritized *GALNT3* from the ADHD+AGG+ASB cross-trait gene-based meta-analysis, because this gene is significantly associated with all three traits and it is nominally associated with thalamus volume (**Supplementary Table 4**).

## Discussion

In this study, we applied two locus-based approaches to localize and characterize shared genetic effects between three genetically correlated traits at the extremes of externalizing behavior: ADHD, AGG, and ASB. We identified eleven significant genes in cross-trait gene-based meta-analyses and 31 shared loci in the local genetic correlation analyses. Through integration of the gene-based and the ρ-HESS-based methodologies, we derived another nine candidate genes that may be involved in the genetic overlap between these traits. Further biological characterization of those 20 prioritized genes highlights a potential role of the amygdala and putamen in externalizing behavior.

Known global genetic correlation estimates for ADHD, AGG, and ASB were obtained from the world-wide largest data sets for each phenotype (Demontis et al., 2019; Pappa et al., 2016; Tielbeek et al., 2017). The high and significant global genetic correlation observed are as reported earlier. While assessing genome-wide genetic covariances between traits at a global level is informative, it does not reflect the actual distribution and size that associations have across regions (Shi et al., 2017). Importantly, at the locus level, the strong positive correlations observed in the global analyses (ADHD+AGG rg = 0.7; ADHD+ASB rg = 0.51) (Hamshere et al., 2013; Zhang-James et al., 2019) were not seen. Although not significant in our study, we observed local heterogeneity of effects when comparing results across different loci in a single cross-trait analysis. Such complex patterns of local genetic covariances have also been described in earlier studies of other psychiatric traits that used the ρ-HESS method (Tylee et al., 2018; Zhu et al., 2019). The co-existence of the large, global positive correlation and the much more heterogeneous ρ-HESS results may be explained by the presence of local, heterogenous pleiotropic effects, especially when the underlying genetic basis involves multiple etiologic pathways. This work illustrates the complex genetic architecture of the overlap between traits and highlights the need for studies to investigate such local, pleiotropic effects.

We identified eleven significant genes in cross-trait gene-based meta-analyses and *GALNT3* showed most pleiotropic effects as it specifically emerged in the three-trait gene-based meta-analysis. One gene was associated with all three traits as identified by the three-trait gene-based meta-analysis of ADHD+AGG+ASB, the polypeptide N-Acetylgalactosaminyltransferase 3 (*GALNT3*). Not much is known about the function of *GALNT3* in relation to brain function and psychiatric disorders, however aberrant glycosylation by N-acetylgalactosaminyl transferases in general has been linked to human cancers and disturbed development and function of the nervous system (Reily, Stewart, Renfrow, & Novak, 2019; Williams, Mealer, Scolnick, Smoller, & Cummings, 2020).

Our combined results from the complementary MAGMA and ρ-HESS analyses identified nine genes. Of those, five genes may be of particular interest because they are highly expressed in the brain and are also nominally associated with brain volumes. Zinc finger protein 521 (encoded by *ZNF521),* a protein acting as transcriptional factor, is involved in the sonic hedgehog pathway; this pathway has an essential role in the central nervous system development (Li et al., 2020; Matsubara et al., 2009; Scicchitano et al., 2019). Genetic variants in *ZNF521* have been associated with irritability (GWAS Atlas; (Watanabe et al., 2019)), which is increasingly recognized as an impairing transdiagnostic symptom underlying multiple internalizing and externalizing disorders (Roy & Comer, 2020). Glycoprotein M6A (coded for by *GPM6A),* is a neural glycoprotein with molecular calcium channel activity, which is expressed in the central nervous system and has been associated with differentiation and migration of neurons derived from stem cells (Michibata et al., 2009; Mukobata et al., 2002; Ramachandran & Margolis, 2017). In addition to its link with neurodevelopment, *GPM6A* also shows significant gene-based association with schizophrenia and educational attainment (GWAS Atlas; (Watanabe et al., 2019)), indicating pleiotropic effects as both lower IQ and behavior problems are associated with increased risk of schizophrenia (Agnew-Blais et al., 2017). Dendrin (*DDN*) is enriched in postsynaptic densities in neurons (Elvira et al., 2006; Herb et al., 1997), however little is known about Dendrin’s function in neurons. *DDN* is associated with a large variety of traits, among others educational attainment and cognitive performance (GWAS Atlas; (Watanabe et al., 2019)) and a previous study suggested that the relation between academic achievement/cognitive ability and externalizing problems may be driven primarily by inattention (Metcalfe, Harvey, & Laws, 2013). Interestingly, *DDN* is also associated with total brain volume (GWAS Atlas; (Watanabe et al., 2019)), and at a genome-wide level ICV (a proxy for total brain volume) negatively correlated with ADHD (Klein et al., 2019). Of note, *DDN* showed nominally significant associations with amygdala and putamen in our analyses. Little is known about the brain-related functions of *ZBTB41* yet, it is highly expressed in brain tissue, and the strongest associations are seen with traits related to factors of the complement system (GWAS Atlas; (Watanabe et al., 2019)), potentially supporting the important role of the complement system in synaptic pruning (Sekar et al., 2016).

Results from the gene expression and gene-based association analyses with brain volumes highlighted several brain regions to be of importance for the specific genes identified and externalizing behaviors in general. The most implicated brain tissues were the cerebellum and frontal cortex (BA9; most of prioritized genes were highly expressed in these tissues) as well as the amygdala and the putamen, for which most of the nominal associations were found. The cerebellum plays a role in the regulation of motor skills, cognitive, and emotional behaviors, and it is known to be involved in neurodevelopmental disorders (Goetz, Schwabova, Hlavka, Ptacek, & Surman, 2017; Stoodley & Limperopoulos, 2016). The frontal cortex region BA9 is important for emotional processing and memory formation (Farrow et al., 2001; Lane et al., 1997). The amygdala is part of the limbic system and an important structure involved emotion processing as amygdala activity has a predominant role in experienced emotional intensity (Frank et al., 2014). The putamen, as part of the dorsal striatum, is involved in stimulus–response habit formation and motor control (Haber, 2016) and a meta-analysis of voxel-based morphometry structural MRI studies of youths with conduct problems showed consistent reductions in grey matter volume of the putamen (Rogers & De Brito, 2016). For all externalizing behaviors studied here, structural changes in cerebellum, frontal cortex, amygdala and putamen have been reported (Fairchild et al., 2019; Hoogman et al., 2017; Johanson, Vaurio, Tiihonen, & Lähteenvuo, 2019; Leutgeb et al., 2015; Moreno-Rius, 2019; Seidman et al., 2006; Waller et al., 2020; Yang & Raine, 2009), which makes these brain regions, and their developmental changes (Teeuw et al., 2018), promising candidates for future studies into the biological basis of the comorbidity of ADHD, AGG, and ASB.

The findings of this study should be viewed in light of strengths and limitations. Instead of following the usual SNP-to-gene approach, we opted for a more robust, region-focused and sensitive approach to study the genetic overlap between ADHD, AGG, and ASB at the locus level. The number of identified loci may seem to be low, and our approach may be limited by the relatively low sample sizes of the AGG, and ASB GWAS datasets. Also, the genetic correlation between AGG and ASB was low, and we cannot rule out a potential sample overlap between both datasets. Efforts to create larger GWAS datasets are ongoing for ADHD, AGG, and ASB, and this opens the possibility of performing better powered analyses, which may be able to provide additional results. Alternatively, phenotypically (and genetically) related traits from large biobank efforts could be informative as well. By integrating results of related traits from the same behavioral domain, such as irritability or other externalizing phenotypes, we may be able to better dissect the underlying biological components. Notably, AGG and ASB can also clearly exists without comorbid ADHD, which means that trait-specific rather than overlapping genetic findings should also be subject of further investigation. Additionally, the biological interpretation of the prioritized genes may be complemented by future studies. Incorporating mRNA expression data of specific neuronal populations (Skene et al., 2018) could be informative to shed more light on the specific functions of the prioritized genes in brain tissues relevant for ADHD, AGG, and ASB. Also, all analyses presented here are based on protein-coding genes. In light of the missing heritability for externalizing behaviors, future studies may want to also integrate information from non-coding genomic segments and other genetic elements, such as structural variants and additional types of rare genetic variation.

In conclusion, this study emphasizes the importance of investigating genetic effects shared between genetically correlated traits at the locus level. We also advocate the use of complementary approaches, since the converging evidence from different cross-trait analysis approaches can highlight additional, novel candidate genes. Uncovering the shared etiological factors underlying externalizing behaviors and related disorders may have important implications for our understanding of the biological mechanisms involved, and ultimately may aid in prevention and/or treatment.

## Acknowledgements

This work was partly carried out on the Dutch national e-infrastructure with the support of SURF Foundation. HHP, BF, and MK are supported by funding for the Dutch National Science Agenda NeurolabNL project (grant 400-17-602). This work was also supported by the European Community’s Horizon 2020 Programme (H2020/2014 – 2020) under grant agreements n° 667302 (CoCA).

## Notes

**Conflict of interest**: Barbara Franke received educational speaking fees from Medice. The other authors declare no conflicts of interests.

### Competing Interest Statement

Barbara Franke received educational speaking fees from Medice. The other authors declare no conflicts of interests.

