## Supplementary Material for "Localizing genomic regions contributing to the extremes of externalizing behavior: ADHD, aggressive and antisocial behaviors"

Supplementary Tables: 4

### Supplementary Tables

**Supplementary Table 1.** Cross-trait genetic correlation analyses result in 31 loci using ρHESS

| chr | start | end | local rg ADHD+AGG | local rg ADHD+ASB |
|---|---|---|---|---|
| 1 | 11777841 | 12779466 | -0.152803121 | 0.420775744 |
| 1 | 59890409 | 61922365 | 0.067999574 | 0.042971589 |
| 1 | 154770403 | 156336133 | 0.483837511 | 0.354359797 |
| 1 | 196176201 | 197311514 | 0.988262402 | -0.960761239 |
| 2 | 110572432 | 113921856 | -0.052472426 | 0.225361764 |
| 3 | 65273538 | 66270447 | 0.035095413 | 0.960313445 |
| 3 | 77508835 | 79024541 | -0.25031436 | 0.422913061 |
| 4 | 6773043 | 7539692 | 0.706325908 | 0.59543495 |
| 4 | 8152235 | 9326479 | -0.045066369 | 0.68953158 |
| 4 | 27343722 | 27965868 | 0.003321683 | -0.157937228 |
| 4 | 97540703 | 99424067 | -0.348038358 | 0.030567005 |
| 4 | 127275477 | 127830293 | -0.465601294 | 0.738451107 |
| 5 | 11940 | 982252 | 0.000550138 | -0.38702438 |
| 5 | 41888710 | 43983499 | -0.335177034 | 0.870562973 |
| 5 | 125785649 | 127344604 | 0.373532123 | -0.003067617 |

| | | | | |
|---|---|---|---|---|
| 6 | 94118142 | 94441175 | 0.688925442 | -0.730273061 |
| 6 | 123856181 | 125424383 | -0.253986859 | 0.618657966 |
| 8 | 11278998 | 13491775 | 0.499317422 | 0.383989982 |
| 8 | 42773823 | 46841315 | 0.884556756 | 0.938747441 |
| 8 | 119685457 | 121201700 | 0.309364949 | 0.105498468 |
| 8 | 133351144 | 134570001 | 0.528742598 | -0.230621225 |
| 10 | 61891409 | 62659679 | -0.647348266 | 0.974585721 |
| 10 | 100668400 | 102949239 | 0.13818094 | 0.551075746 |
| 10 | 116421406 | 119523934 | 0.940949363 | -0.547744192 |
| 11 | 89208936 | 90966490 | -0.689860672 | -0.647230002 |
| 12 | 124977980 | 126445505 | -0.125739598 | -0.533918054 |
| 13 | 104066710 | 104844114 | -0.138924264 | -0.862517486 |
| 14 | 28466533 | 29972145 | -0.721808643 | 0.299104259 |
| 17 | 36809344 | 38877404 | 0.317404181 | -0.035964118 |
| 18 | 26875587 | 27866478 | 0.003438467 | 0.615754626 |
| 22 | 37570269 | 39307894 | 0.295994237 | 0.026228558 |

chr = Chromosome, start = locus start position (in hg19), end = locus end position (in hg 19), rg = genetic correlation.

**Supplementary Table 2**. Results of single- and cross-trait gene-based meta-analysis association analyses for all genes identified by combining ρHESS and MAGMA analyses.

| GENE | ADHD+AGG MA GENE-BASED P | ADHD+ASB MA GENE-BASED P | ADHD GENE-BASED P | AGG GENE-BASED P | ASB GENE-BASED P |
|---|---|---|---|---|---|
| ***PBXIP1*** | 0.0033 | 0.017 | 0.0088 | 0.093 | 0.477 |
| ***PYGO2*** | **0.0006** | **0.0004** | **0.0006** | NA | 0.126 |
| ***KCNT2*** | 0.036 | 0.021 | 0.060 | 0.185 | 0.084 |
| ***CFHR1*** | 0.038 | 0.043 | 0.052 | 0.233 | 0.273 |
| ***F13B*** | 0.034 | 0.025 | 0.198 | 0.014 | 0.0055 |
| ***ZBTB41*** | 0.0074 | **0.0010** | 0.037 | 0.037 | **0.0007** |
| ***CHUK*** | 0.040 | 0.044 | 0.011 | 0.690 | 0.745 |
| ***CWF19L1*** | 0.013 | 0.023 | 0.0038 | 0.566 | 0.771 |

| | | | | | |
|---|---|---|---|---|---|
| ***BLOC1S2*** | 0.011 | 0.045 | 0.0072 | 0.360 | 0.829 |
| ***ATRNL1*** | **0.00002** | **0.0002** | **0.0001** | 0.030 | 0.214 |
| ***KCNK18*** | 0.037 | 0.0052 | 0.043 | 0.278 | 0.014 |
| ***TMEM132B*** | 0.0019 | 0.0026 | 0.0038 | 0.122 | 0.172 |
| ***MLLT6*** | 0.031 | 0.041 | 0.021 | 0.419 | 0.542 |
| ***PCGF2*** | 0.017 | 0.0031 | 0.0063 | 0.547 | 0.127 |
| ***ZC3H8*** | 0.014 | 0.026 | 0.0057 | 0.504 | 0.724 |
| ***ZC3H6*** | 0.012 | 0.0045 | 0.010 | 0.318 | 0.117 |
| ***TTL*** | 0.025 | 0.0022 | 0.012 | 0.507 | 0.038 |
| ***ANKRD54*** | 0.025 | 0.048 | 0.0067 | 0.653 | 0.855 |
| ***TMEM184B*** | 0.0044 | 0.029 | 0.0069 | 0.168 | 0.718 |
| ***GTPBP1*** | 0.042 | 0.021 | 0.030 | 0.430 | 0.215 |
| ***SUN2*** | 0.026 | 0.011 | 0.012 | 0.516 | 0.263 |
| ***TBC1D14*** | 0.0023 | **0.0003** | 0.0053 | 0.108 | 0.0077 |
| ***C5ORF51*** | **0.0004** | 0.022 | 0.0055 | 0.0087 | 0.675 |
| ***FBXO4*** | **0.0003** | 0.0097 | **0.0014** | 0.048 | 0.720 |
| ***GHR*** | 0.0077 | 0.0029 | 0.0032 | 0.454 | 0.219 |
| ***CCDC152*** | **0.0014** | 0.0034 | 0.0098 | 0.026 | 0.085 |
| ***SEPP1*** | **0.0009** | 0.012 | 0.014 | 0.0069 | 0.237 |
| ***ZNF131*** | 0.018 | 0.043 | 0.020 | 0.266 | 0.575 |
| ***HMGCS1*** | 0.032 | 0.0090 | 0.076 | 0.108 | 0.010 |
| ***NKAIN2*** | 0.012 | **0.00147** | 0.015 | 0.227 | 0.013 |
| ***GATA4*** | 0.0032 | 0.0091 | 0.0050 | 0.160 | 0.417 |
| ***TMEM71*** | 0.021 | 0.044 | 0.012 | 0.438 | 0.719 |
| ***TG*** | 0.0033 | 0.011 | 0.0095 | 0.086 | 0.321 |
| ***SLA*** | 0.035 | 0.048 | 0.134 | 0.042 | 0.073 |

Gene-based association results are shown for all 34 genes overlapping between ρHESS and MAGMA cross-trait analyses (**Figure 1, Step C**). Significant gene p-values ($p < 0.05/34$) are marked in bold. P = p-value; MA = cross-trait gene-based meta-analysis.

**Supplementary Table 3**. GTEx median expression values of top 5% tissues for prioritized genes

| Genes | Top 5% tissues | TPM |
|---|---|---|
| **ADHD+AGG MA** | | |
| ***ZNF521*** | **Brain; Cerebellar Hemisphere** | 104.9 |
| | **Brain; Cerebellum** | 83.33 |
| | Cervix; Ectocervix | 21.97 |
| ***MECOM*** | Stomach | 33.88 |
| | Kidney; Medulla | 23.41 |
| | Lung | 15.58 |
| **ADHD+ASB MA** | | |
| ***HYI*** | NA | NA |
| ***DDN*** | **Brain; Anterior cingulate cortex (BA24)** | 301 |
| | **Brain; Frontal Cortex (BA9)** | 268.1 |
| | **Brain; Nucleus accumbens (basal ganglia)** | 266.8 |
| ***FAM186B*** | Testis | 60.94 |
| | Ovary | 1.063 |
| | Thyroid | 0.9144 |
| ***GPM6A*** | **Brain; Cerebellar Hemisphere** | 460.7 |
| | **Brain; Frontal Cortex (BA9)** | 335.2 |
| | **Brain; Anterior cingulate cortex (BA24)** | 285.6 |
| ***COL19A1*** | Cells; EBV-transformed lymphocytes | 6.549 |
| | **Brain; Cerebellum** | 2.943 |
| | Bladder | 2.429 |
| ***LY96*** | Spleen | 73.37 |
| | Cells; Cultured fibroblasts | 32.97 |
| | Cells; EBV-transformed lymphocytes | 31.8 |
| ***SZT2*** | Thyroid | 25.25 |
| | Testis | 24.84 |
| | Ovary | 22.76 |
| ***DUSP6*** | Adipose; Visceral (Omentum) | 91.55 |
| | Spleen | 81.47 |

| | | |
|---|---|---|
| | Lung | 74.64 |
| **ADHD+AGG+ASB MA** | | |
| ***GALNT3*** | Minor Salivary Gland | 31 |
| | Esophagus; Mucosa | 25.44 |
| | Stomach | 19.08 |
| **p-HESS+MAGMA** | | |
| ***PYGO2*** | Pituitary | 65.53 |
| | Thyroid | 61.63 |
| | Cervix; Endocervix | 58.63 |
| ***ZBTB41*** | **Brain - Cerebellar Hemisphere** | 20.44 |
| | **Brain - Cerebellum** | 19.07 |
| | Cells - Cultured fibroblasts | 9.431 |
| ***ATRNL1*** | **Brain; Frontal Cortex (BA9)** | 24.84 |
| | **Brain; Cortex** | 17.15 |
| | **Brain; Anterior cingulate cortex (BA24)** | 10.34 |
| ***TBC1D14*** | Adrenal Gland | 23.68 |
| | Testis | 23.61 |
| | Stomach | 23.49 |
| ***C5orf51*** | Cells; Cultured fibroblasts | 34.08 |
| | **Brain; Cerebellar Hemisphere** | 22.63 |
| | Cells; EBV-transformed lymphocytes | 20.64 |
| ***FBXO4*** | Adipose; Subcutaneous | 13.03 |
| | Nerve; Tibial | 12.67 |
| | Cells; Cultured fibroblasts | 12.29 |
| ***CCDC152*** | Cells; EBV-transformed lymphocytes | 10.62 |
| | Ovary | 10.54 |
| | Nerve; Tibial | 10.47 |
| ***SEPP1*** | Brain; Spinal cord (cervical c-1) | 331.7 |
| | Cervix; Ectocervix | 222.1 |
| | Ovary | 217.6 |
| ***NKAIN2*** | Brain; Spinal cord (cervical c-1) | 85.98 |

| | | |
|---|---|---|
| | **Brain; Substantia nigra** | 23.25 |
| | **Brain; Hippocampus** | 21.85 |

Data from the GTEx project (https://gtexportal.org/home/) are shown for the 20 genes prioritized by the different approaches. Results show the tissues within the top 5% tissues (3 out of 54) that most strongly express a gene (row) based on the median Transcripts Per Million (TPM). Tissues in bold mark the 12 brain tissues. No data was available for the *HYI* gene. NA = Not available.

**Supplementary Table 4**. Results of gene-based association analyses with (subcortical) brain volumes.

| GENE | ACCUMBENS P | AMYGDALA P | CAUDATE P | HIPPOCAMPUS P | ICV P | PALLIDUM P | PUTAMEN P | THALAMUS P |
|---|---|---|---|---|---|---|---|---|
| | | | | **ADHD+AGG** | | | | |
| ***ZNF521*** | 0.876 | 0.337 | 0.063 | 0.012289* | 0.704 | 0.472 | 0.0026811* | 0.135 |
| ***MECOM*** | 0.112 | 0.021873* | 0.816 | 0.911 | 0.541 | 0.035301* | 0.169 | 0.0054397* |
| | | | | **ADHD+ASB** | | | | |
| ***HYI*** | 0.179 | 0.246 | 0.300 | 0.574 | 0.729 | 0.255 | 0.295 | 0.254 |
| ***DDN*** | 0.384 | 0.036758* | 0.834 | 0.469 | 0.861 | 0.785 | 0.0016475* | 0.397 |
| ***FAM186B*** | 0.436 | 0.682 | 0.897 | 0.213 | 0.245 | 0.143 | 0.137 | 0.476 |
| ***GPM6A*** | 0.460 | 0.945 | 0.429 | 0.917 | 0.329 | 0.952 | 0.006155* | 0.769 |
| ***COL19A1*** | 0.637 | 0.542 | 0.211 | 0.558 | 0.268 | 0.458 | 0.157 | 0.242 |
| ***LY96*** | 0.363 | 0.869 | 0.391 | 0.474 | 0.484 | 0.324 | 0.084 | 0.064 |
| ***SZT2*** | 0.047467* | 0.869 | 0.305 | 0.645 | 0.628 | 0.131 | 0.441 | 0.079 |
| ***DUSP6*** | 0.686 | 0.331 | 0.0015934* | 0.475 | 0.490 | 0.342 | 0.912 | 0.886 |
| | | | | **ADHD+AGG+ASB** | | | | |
| ***GALNT3*** | 0.653 | 0.905 | 0.733 | 0.082 | 0.149 | 0.447 | 0.077 | 0.018297* |
| | | | | **PHESS+MAGMA** | | | | |
| ***PYGO2*** | 0.313 | 0.279 | 0.696 | 0.038036* | 0.199 | 0.638 | 0.030397* | 0.699 |
| ***ZBTB41*** | 0.293 | 0.747 | 0.092 | 0.906 | 0.440 | 0.014296* | 0.170 | 0.157 |
| ***ATRNL1*** | 0.083 | 0.546 | 0.792 | 0.421 | 0.720 | 0.324 | 0.484 | 0.676 |
| ***TBC1D14*** | 0.039184* | 0.841 | 0.224 | 0.687 | 0.576 | 0.145 | 0.277 | 0.209 |
| ***C5ORF51*** | 0.900 | 0.696 | 0.876 | 0.387 | 0.372 | 0.580 | 0.812 | 0.589 |
| ***FBXO4*** | 0.827 | 0.734 | 0.858 | 0.215 | 0.389 | 0.740 | 0.674 | 0.564 |

| | | | | | | | | |
|---|---|---|---|---|---|---|---|---|
| ***CCDC152*** | 0.318 | 0.036989* | 0.802 | 0.751 | 0.216 | 0.702 | 0.199 | 0.233 |
| ***SEPP1*** | 0.409 | 0.028106* | 0.874 | 0.841 | 0.189 | 0.564 | 0.195 | 0.415 |
| ***NKAIN2*** | 0.862 | 0.921 | 0.810 | 0.948 | 0.111 | 0.772 | 0.991 | 0.466 |

Gene-based association results are shown for the seven subcortical brain volumes and ICV (Adams et al., 2016; Hibar et al., 2017; Satizabal et al., 2019). In total, 20 prioritized genes from the different approaches were tested. None of the genes was significantly associated after correction for multiple testing (p-value < 0.05/(20*8)). *indicates nominally significant gene-based association (p < 0.05). ICV = Intracranial volume; P = p-value.
